# Silver nanoparticles can be sampled by ultrafiltration probe but elution into & recovery from plasma and DPBS differs *in vitro*

**DOI:** 10.1101/2024.01.26.577516

**Authors:** Marije Risselada, Robyn R McCain, Miriam G Bates, Makensie L Anderson

**Affiliations:** Department of Veterinary Clinical Sciences, College of Veterinary Medicine, Purdue University, West Lafayette, Indiana, United States of America; Center for Clinical Translational Research, College of Veterinary Medicine, Purdue University, West Lafayette, Indiana, United States of America

## Abstract

We compared 1) the influence of elution fluid on rate, pattern, and completeness of silver nanoparticle (AgNP) elution, and 2) ultrafiltration (UF) probe and direct sampling *in vitro*. Six specimens (2.5ml of 0.02mg/ml 10nm AgNP and 5.0ml of 30% poloxamer 407) contained in a dialysis tube (12-14kDa pores) were placed in 100ml Dulbecco’s Phosphate Buffered Saline (DPBS) (n=3) or canine plasma (n=3) for 96h on a stirred hot plate (37°C and 600rpm) and sampled 20 times. Six pipette and UF probe samples were taken of a 0.001mg AgNP/ml DPBS or plasma solution. Inductively coupled plasma mass spectrometry was used to analyze Ag. Stock plasma contained Ag. At 96h, 5/6 dialysis tubes had not fully released AgNP. One peak in hourly Ag increase was present in DPBS (10-13h), and two peaks in plasma (6-8h and 10-13h). The hourly Ag increase in plasma decreased earlier than in DPBS. UF probe sampling was possible in both DPBS and plasma and resulted in higher Ag concentrations but with more variation than pipette samples. While *in vitro* use of DPBS might be more cost effective, plasma should be considered due to difference in elution and recovery.

## Introduction

With the increase in antimicrobial resistance in veterinary medicine [1,2], novel strategies and antimicrobials to combat infections are (re)gaining favor. One strategy might be to deliver a high local dose of antibiotics [3–7]. Another might be to explore the use of non-antibiotic antimicrobials, such as silver (Ag), especially silver nanoparticles (AgNP). They have received considerable interest for their use against micro-organisms [8,9] and in wound care [10]. Poloxamer 407 is a versatile reverse gelatinating polymer that is safe for implantable use [11,12], and has found use for delivery of antifungals [12], chemotherapeutics [11,13], and antibiotics [14]. It also might have inherent antimicrobial properties of its own [15]. Prior silver elution studies have been performed into phosphate-buffered saline (PBS) from poloxamer 407 [16], into deionized water baths (from silver oxide films) [17], or aqueous samples with varying NaCl concentrations (from solid coated specimens) [18]. Plasma might be a closer *in vitro* fluid approximation to an *in vivo* environment than PBS and could mimic protein interactions that might be encountered. The elution into human plasma from an antibiotic containing polymethyl methacrylate (PMMA) construct as well as the effect plasma on PMMA properties was reported [19]. However, plasma has not been evaluated for suitability of Ag release studies, and whether the benefit could outweigh the increased cost of commercial canine plasma (∼$2,400/500ml; Innovative research, Novi, MI) over PBS (∼$30/500ml; Gibco, Thermofisher Scientific, Billings, MT).

The ultimate goal of sustained release compounds assessed by elution studies is their *in vivo* application. Ultrafiltration (UF) probes have been used successfully in dogs [20,21] and are used to sample compounds present in the tissue, without the need for more invasive sampling methods, such as tissue biopsies [22]. Establishing the feasibility of sampling Ag with UF probes would be preferable prior to their use to investigate the *in vivo* pharmacokinetic profile of Ag containing local delivery products. However, the possibility of sampling AgNP via UF probe has not been assessed, nor has a comparison of canine UF probes and direct sampling been reported for AgNP.

We aimed to 1) determine and compare the influence of elution fluid on rate, pattern, and completeness of AgNP elution *in vitro*, and 2) compare UF probe sampling with direct samples. We hypothesized that 1) elution of AgNP into Dulbecco’s PBS (DPBS) will be similar to elution in canine plasma, and that 2) UF probe sampling will be constant over time and yield similar results as direct sampling.

## Materials and Methods

An observational elution study and repeat sampling study were performed separately. Canine Ultrafiltration Probes (BASi Instruments, West-Lafayette, IN) were assembled as per manufacturer instructions [23].

### Elution

A commercial 0.02mg/ml 10nm silver nanoparticles (AgNP) in aqueous buffer with sodium citrate as stabilizer (Sigma Aldrich, Saint Louis, MO) was used for the study. Six elution specimens were prepared. Each contained 2.5ml of the 0.02 mg/ml 10nm AgNP stock solution mixed with 5.0ml 30% poloxamer 407 (Pluronic® F-127, Sigma Aldrich) (1:2 ratio with a total Ag content of 0.05mg/specimen). The specimens were prepared in individual 12ml syringes <2 hours before use and stored refrigerated and shielded from light. Individual 150ml crystallization dishes (Synthware glass, Pleasant Prairie, WI) with 100ml fluid were prewarmed for 2 hours prior to starting the elution study. Three dishes had Dulbecco’s phosphate-buffered saline without Calcium chloride or magnesium chloride added (DPBS, Gibco, Thermofisher Scientific, Billings, MT) and three had canine plasma with dipotassium EDTA (K2-EDTA) (IGCNPLAK2E500ml Canine Plasma lot 41273, Innovative research, Novi, MI). A 10 multi-position hotplate with magnetic stir bars was used (RT 10, IKA Magnetic Stirrers, Wilmington, NC) with settings at 37°C and 600 rpm throughout the experiment. Room temperature was set at 68°F. Specimens were created using a 10cm long strip of 1 inch dialysis tubing with 12-14kDa pores (Carolina Biological Supply Co, Burlington, NC) [24]). The distal free end was folded up length wise and secured in folded position using 2 large surgical stainless steel hemoclips (Hemoclips® Teleflex, Morrisville, NC) in opposite direction [16]. The 1:2 AgNP-poloxamer mix was then placed in the tube, and the proximal free end of the tube folded and closed in similar fashion. All specimens were created in one batch and then placed in the prewarmed DPBS or canine plasma within 5 minutes after assembly. The t=0 sample was taken immediately after all specimens were submerged in the same order of assembly and placement. Twenty samples (0.15ml each) were taken over 96 hours with a decrease in frequency (0, 1, 2, 3, 4, 5, 6, 8, 10, 13, 17, 22, 27, 34, 42, 48, 58, 66, 72, 96 hours) with the sampling order consistent throughout. At 96 hours, an additional sample (by needle aspiration through the tube) of the fluid contained within the dialysis tube was taken.

### Ultrafiltration probe sampling

Two specimens were prepared immediately prior to sampling in a 50ml conical centrifuge tube (Thermofisher Scientific) using graduated pipettes (Thermofisher Scientific) and a 3ml syringe (Covidien, Mansfield, MA). Each specimen contained 0.03mg of AgNP (1.5ml of the commercial 0.02mg AgNP stock solution) total in 28.5ml of either DPBS (n=1) or canine plasma (n=1) for a targeted fluid concentration of 0.001mg/ml of AgNP. Ultrafiltration probes were tested for patency and sampling using DPBS prior to use for the study.

Full submerging of the probe near the lowest point of the tube was ensured, and positioning was checked after each manipulation. Vacutainers without additives (Vacuette Blood collection tube, 3.0ml, no additive, Greiner Bio-One, Sigma Aldrich, St Louis, MO) were used to collect the probe samples and an additional 10ml of air was removed to increase negative pressure in each vacutainer to increase sampling speed. The initial sample of the study specimen was discarded to avoid risk of dilution by plain DPBS. Repeat samples using both a UF probe and direct sampling using a pipette from the same area as the membrane of the UF probe were taken at 0, 5, 10, 15, 30 and 60 minutes from the plasma and DPBS specimen.

Samples of commercial stock AgNP solution, DPBS and canine plasma were taken prior to the start of the study. All samples were stored at −78°C until batch analysis.

### Sample and data analysis

The quantity of Ag in each sample was determined via inductively coupled plasma mass spectrometry (ICP-MS) as previously described [16]. DPBS samples were diluted in 2% HNO3 and plasma samples were digested over night at 70°C in the 1:1 mixture of 70% HNO3 and H2O2 and analyzed using ICP-MS (Perkin Elmer NexION 300D) to determine the concentration of silver within each sample. The short-term precision was less than 3% relative standard deviation (RSD), and the long-term stability was <4% relative standard deviation over 4 hours. Isotope-ratio precision was less than 0.08% relative standard deviation. The Ag detection limit was 0.001ng/ml, and quantification limit at 0.002ng/ml, and all samples below this limit were recorded as 0ng/ml. Silver concentrations were expressed in parts per billion (ppb), with 1ppb=1ng/ml. Hourly increase in Ag was calculated as the difference between the measured values at two subsequent time points divided by the hours between time points (expressed as ppb/hr). The data of the 3 elution specimens will be expressed graphically as mean±SD. The values of the repeat sampling experiment will be reported as individual results and a mean±SD (RSD) for both specimens in an observational manner without further statistical analysis. The RSD was calculated by dividing the SD by the mean and will be expressed as a %.

## Results

Silver concentrations for stock solutions used in this study were: DPBS stock 0.19ppb Ag; plasma stock 4.83ppb Ag and the commercial 0.02mg/ml AgNP solution contained 24,940ppb Ag. Assembled UF probes initially did not reliably yield appropriate negative suction to obtain a sample, and probes were not re-usable for repeat experiments. Additional negative pressure applied to the vacutainers together with sealing connecting points with glue (3g Tube, The Gorilla Glue Company, Cincinnati, OH) allowed single use sampling.

### Elution

No leakage of any dialysis tubes was observed under gentle pressure immediately after assembly. Five out of 6 dialysis tubes were fully filled at 96 hours, the sixth was not fully filled when retrieved. The five fully filled specimens still contained more Ag than the surrounding fluid at 96 hours (Table 1), and release of Ag was not complete at 96 hours. A burst release of Ag was seen both into DPBS and plasma in the first 13 hours (DPBS, light grey) and 8 hours (plasma, dark grey), with the baseline amount in plasma higher than in DPBS (Fig 1AB). The increase of Ag measured in plasma decreased earlier, between 8-60 hours (dark grey, Fig 1AB), and the measured Ag decreased thereafter. The hourly increase of Ag in DPBS (light grey) tapered between 27-84 hours but continued throughout the study (Fig 1C). The highest hourly increase in Ag was between 10-13 hours in both DPBS (light grey) and plasma (dark grey) (Fig 1D). A clear single peak in DPBS (20.66ppb/hr) whereas two peaks were present for plasma: between 6-8 hours (12.31ppb/hr) and between 10-13 hours (13.85ppb/hr) (Fig 1D). The amount of Ag measured at 96 hours was higher in the remaining Ag:poloxamer mix contained within the tube than the surrounding fluid (both DPBS and plasma) (Fig 2). The Ag concentration measured at 96 hours in DPBS was higher (both specimen and fluid) than in plasma (Fig 2). The mean±SD total amount of Ag removed via sampling for the three DPBS fluid set ups was 330±51ng Ag and for the three plasma fluid set ups 198.1±71ng Ag.

**Fig 1:**
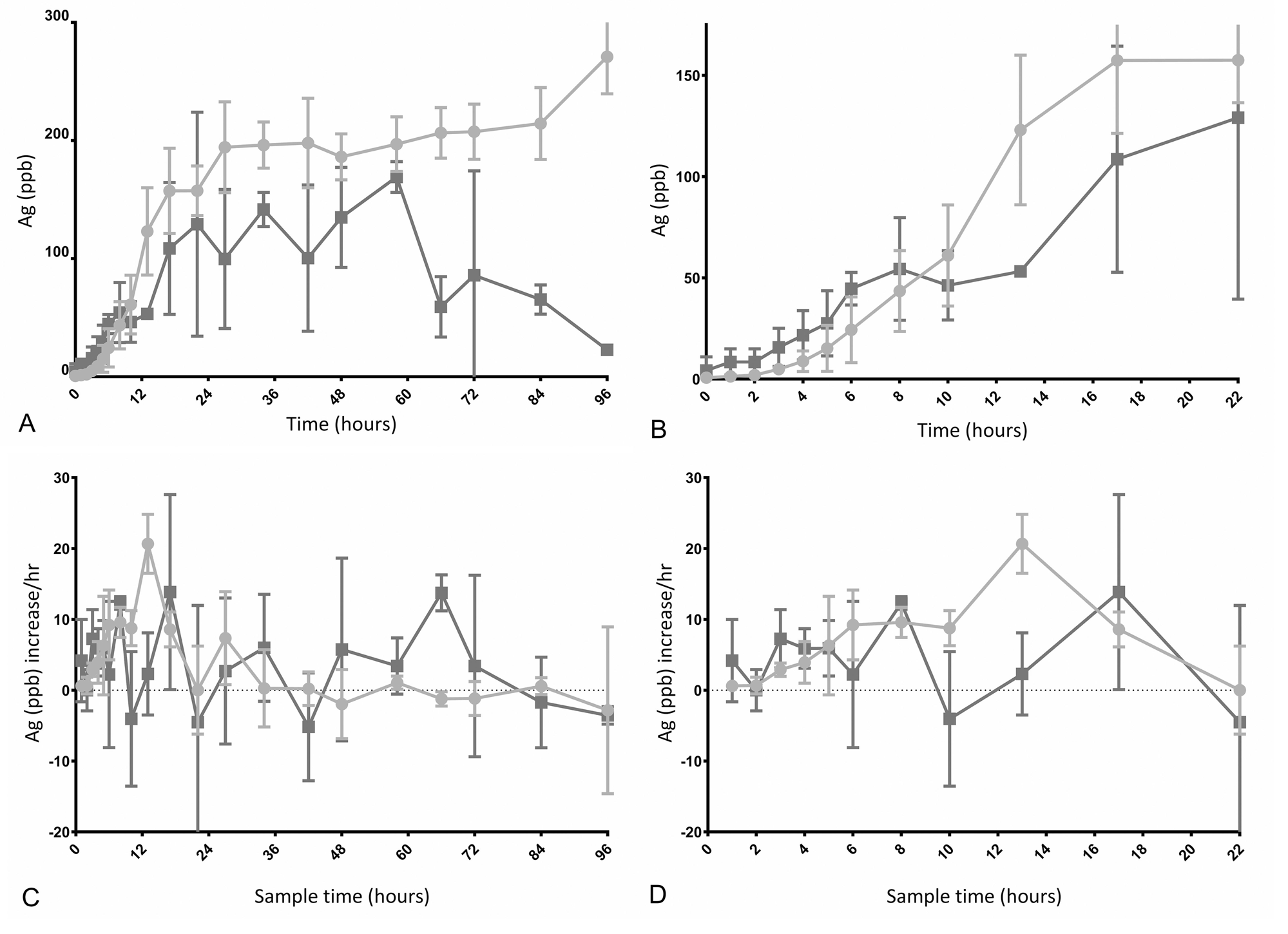
A. Elution of Ag into either DPBS (light grey) or plasma (dark grey), expressed as ppb Ag over 96 hours. The elution of Ag into DBPS continued longer, leading to a higher fluid concentration, whereas the increase of Ag in plasma tapered between 24 - 60 hours and Ag decreased after 60 hours. B. The increase in Ag is expressed as Ag (ppb)/hour measured over the timeframe prior to the sampling time. A burst elution pattern is evident for both elution fluids. C. Elution of Ag into either DPBS or plasma, expressed as ppb Ag over the first 22 hours. D. Hourly increase in Ag over the first 22 hours. The measured values of Ag in plasma varied more for each time point and between time points as evidenced by the larger bars and the shape of the curve.

**Fig 2:**
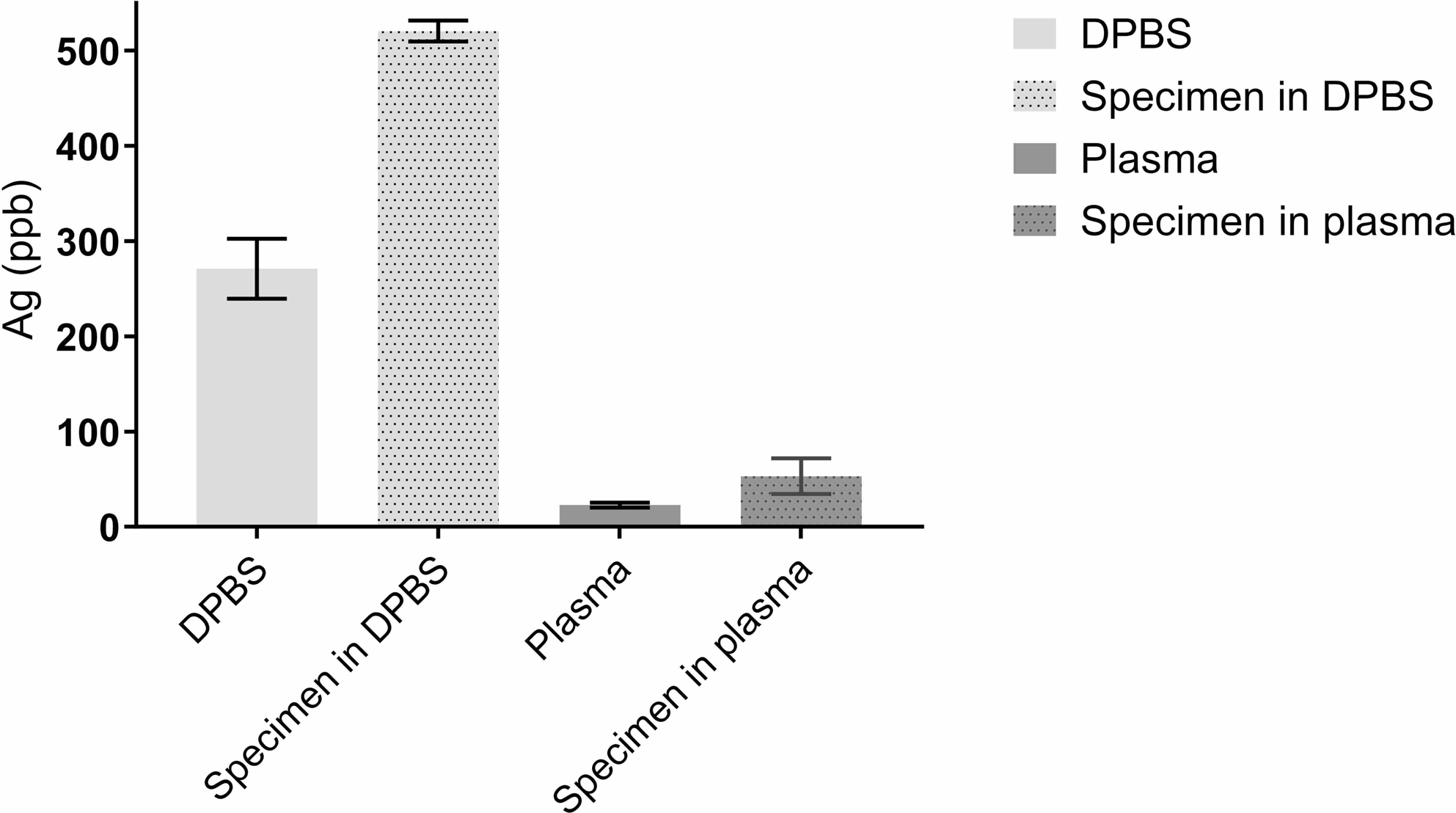
Ag amount (ppb) in elution fluid (DPBS or plasma) and remainder of the Ag:poloxamer specimen at 96 hours. DPBS elution fluid is shown in light grey, with the remaining specimen in shaded light grey. The plasma elution fluid is shown in dark gray. Measured Ag content in both the fluid and Ag:poloxamer specimen remnant at 96 hours were less in plasma than DPBS, with the specimens still higher in Ag content than the surrounding DPBS or plasma.

**Table 1:**
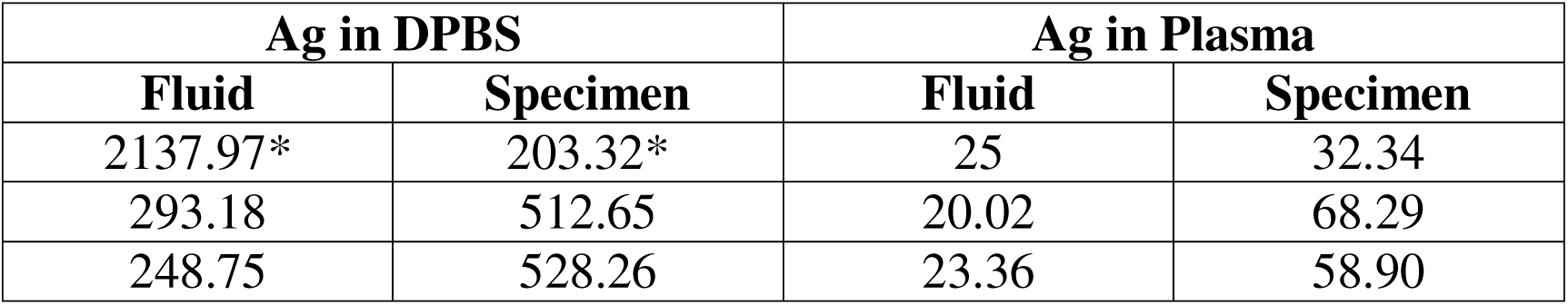
Silver (Ag) at 96 hours expressed as parts per billion (ppb) for the remaining elution fluid outside of the dialysis tube and the remaining specimen contained within the dialysis tube. Each tube contained 6660ppb Ag at the start of the experiment. *denotes the specimen that was not fully filled and the surrounding elution fluid. Silver at 96 hours was analyzed once for six specimens (3 each in DPBS and plasma) and elution fluid.

### Ultrafiltration probe sampling

Anticipated Ag content based on the stock (24,940ppb Ag (AgNP stock) diluted in 30ml) was 1,247ppb Ag of the diluted fluid. Ultrafiltration probes were able to collect Ag in both DPBS and plasma, with Ag content in fluid obtained via UF probe sampling higher than the corresponding pipette samples (Fig 3). However, the UF probe-obtained samples had more variation and values differed between sampling times and decreased over time (Table 2). Samples obtained by both methods had a measured Ag lower than anticipated, with the underestimation greater in DPBS than plasma.

**Fig 3:**
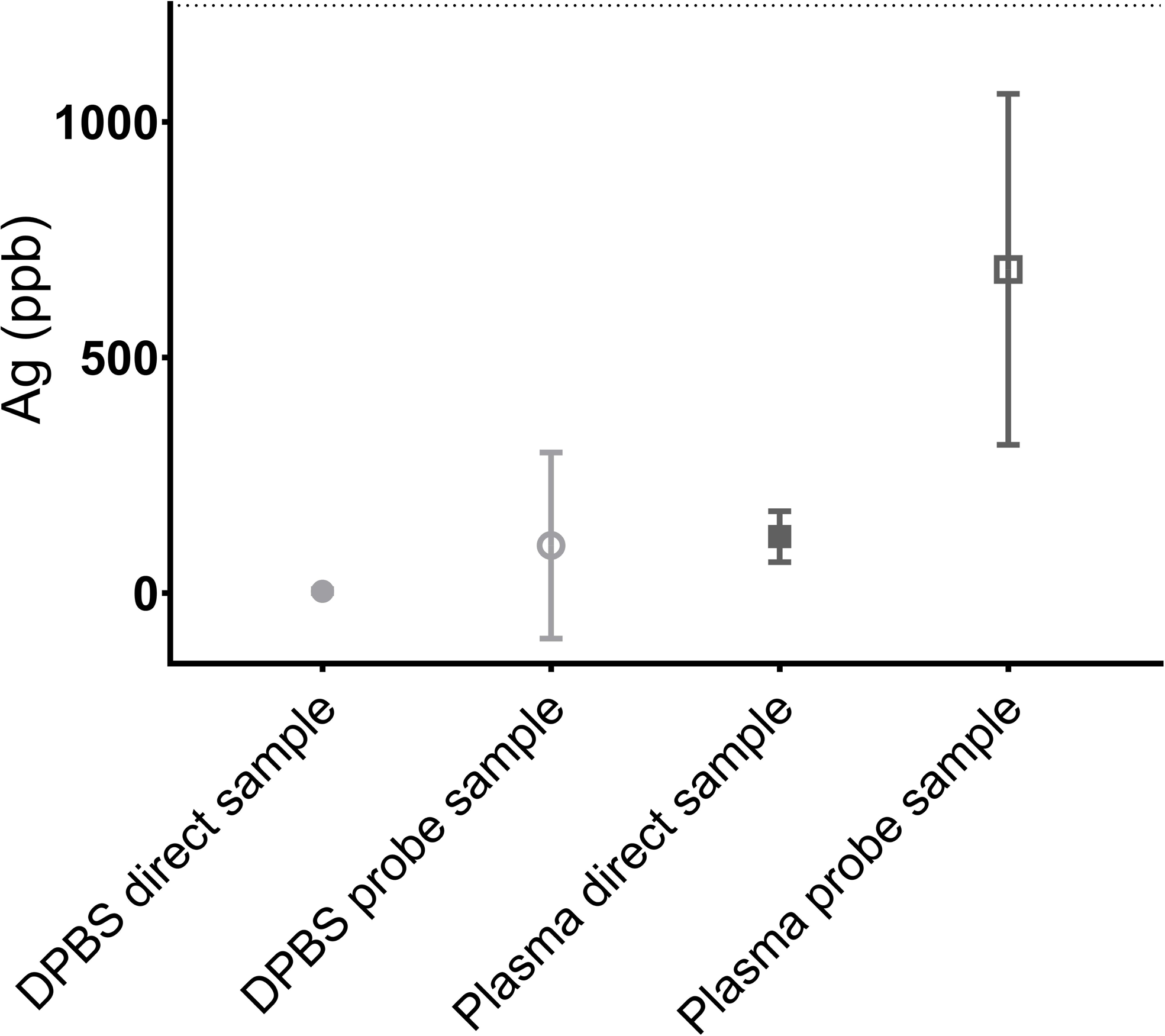
Pipette and ultrafiltration (UF) probe sampling of 1,247ppb Ag in DPBS (light grey) and plasma (dark grey) solution. Pipette sampling is represented by a solid sphere (DPBS) or square (plasma) symbol, while the corresponding UF sampling data is shown by a solid sphere or square. Both methods underestimated the Ag content in both mediums. The samples obtained via UF probe had a wider variation in measured Ag. The dotted line indicates 1,247ppb Ag.

**Table 2:**
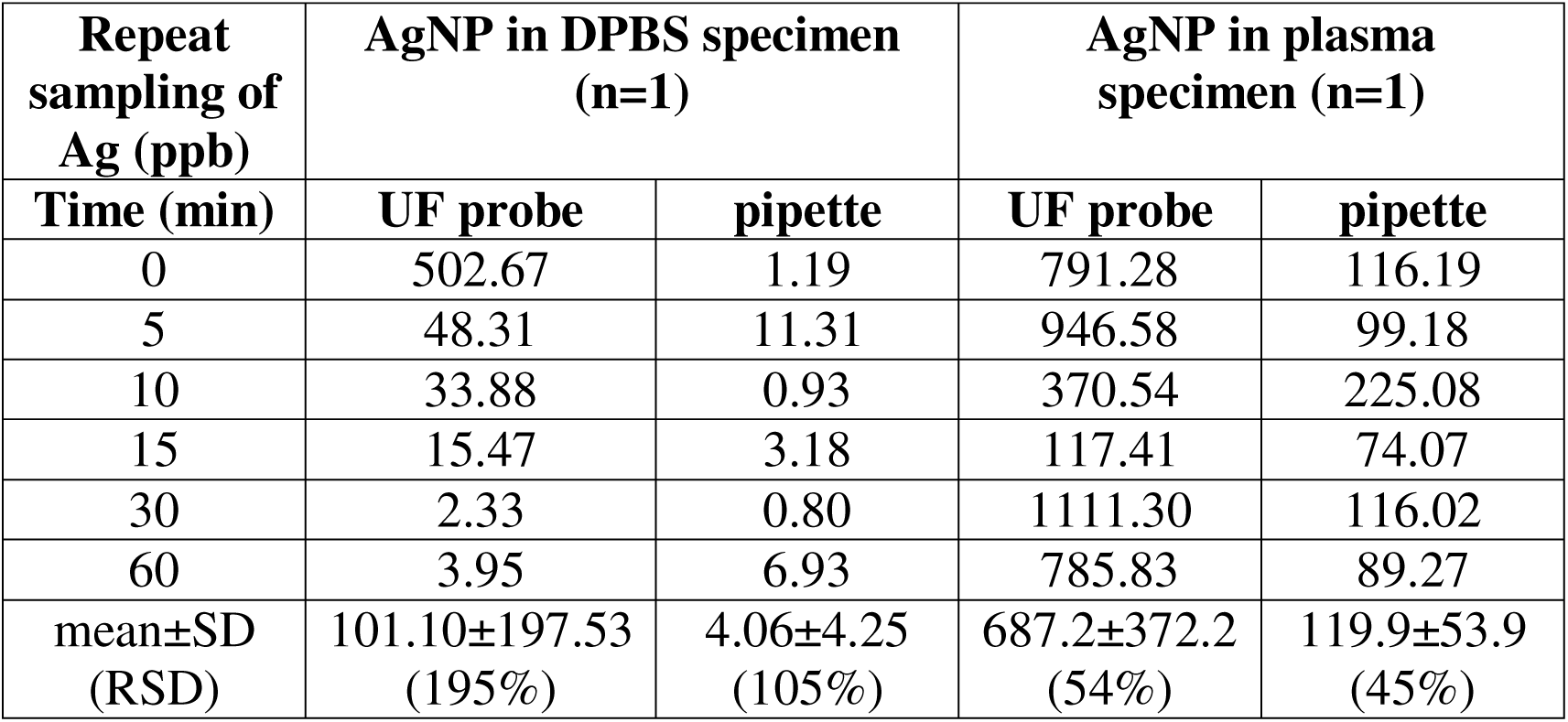
Repeat pipette and ultrafiltration (UF) probe sampling of a planned solution of 1,247ppb Ag in DPBS and plasma solution (one specimen each). Probes were allowed to sample for 4 minutes to obtain enough sample volume. Direct samples were taken using a pipettor at the time point. UF probe samples were taken by attaching a vacutainer to the probe for 4 minutes at the start time. DPBS= Dulbecco’s Phosphate Buffered Saline, UF= ultrafiltration. The mean is provided with both standard deviation (SD) and relative standard deviation (RSD) for each sampling method over time from the same fluid.

## Discussion

Elution into DPBS yielded different results than elution into plasma, with a longer sustained initial hourly increase in fluid Ag and more variation in Ag measurements with a resultant less smooth curve. A burst release pattern was evident for both DBPS and plasma as elution fluid. The burst release corresponds with an earlier study where ∼88% of the Ag was released from poloxamer 407 within the first 24 hours [16]. While the pore size of the dialysis tube should allow AgNPs to cross freely, the Ag concentration within the dialysis tube was still higher at 96 hours than the surrounding fluid (both DPBS and plasma), indicating incomplete release of the specimen in both elution media. Silver could be measured in DPBS and plasma by UF probe sampling.

The higher initial values of Ag found in plasma might be explained by Ag being present in stock plasma while no Ag was found in the DPBS stock solution. The higher initial value in plasma was then followed by an increase in Ag content similar in shape to the DPBS curve.

However, the presence of Ag, and the initial starting value, would mask the initial release of Ag at the time when the release will be highest and is a limitation of using plasma as the elution fluid. We chose to report the measured values as-is instead of detracting the initial Ag measured obtained from stock plasma, as we only had a single measurement for the stock plasma rather than repeat samples and thus felt that correcting could inadvertently introduce error as well.

Plasma with EDTA was chosen as it was the cheapest commercial option available. Ethylenediaminetetraacetic acid (EDTA) chelates silver [25], and its presence in plasma theoretically could help extract Ag from the specimen contained within the dialysis tube. However, chelation of Ag into stable complexes also might hinder analyses, although measuring of excreted EDTA-metal complexes before and during chelation therapy has been described [25,26]. In addition, nitric acid digestion has been used to separate heavy metals from EDTA [27]. However, depending on the compound for which the elution is performed and its chelation abilities with EDTA plasma with a different anti-coagulation agent might be preferred.

We used a 1:2 ratio of AgNP:poloxamer in this study to maximize the Ag content while still maintaining the full gelatinating properties of the poloxamer. In a prior elution study, a 1:4 ratio of AgNP:poloxamer was used and the specimens fully gelatinated [4]. However, specimens did not fully gel in a 1:1 ratio in a different study [15] but did when we tested at a 1:2 ratio (unpublished data). The specimen composition, as well as methodology (a single, larger amount of elution fluid with sampling as opposed to full exchange of smaller amounts of elution fluid) between this study and the prior study might account for the earlier tapering of elution and the flattening of the curve. The fluid exchange would increase the gradient and therefore drive migration of Ag across the dialysis tube membrane. The fluid quantity was chosen to allow a model with a larger amount of fluid without full exchange to allow continuous stirring, the ability to submerge a large specimen, and due to the cost of canine plasma (∼$2,400 per 500ml). Removing a sample at each time point (0.15ml each for a total of 3 ml) could impact the Ag concentration by both removing fluid as well as Ag. The total amount of Ag removed over the entirety of the 96 hours was 330ng for the DPBS fluid set ups and 158ng for the plasma set ups. Ideally the removed amount of Ag would have been added back in a corrected calculating using the currently present volume of the elution fluid, however, the fluid present at each time point was not measured, and using an approximation might lead to a flawed correction, and more error than not correcting.

The dialysis tube model was chosen to avoid the poloxamer from dissolving immediately upon placing the AgNP:poloxamer mix in the elution DPBS or plasma fluid. Prior studies with the same commercial AgNP and dialysis tubing [16] yielded appropriate migration of Ag through the membrane. Given that no plasma was placed within the tube, there were no concerns of Ag complexing with proteins, leading to inability to migrate across the membrane due to pore size (12-14kDa).

While aluminum foil was wrapped tightly around the opening of the dishes, fluid loss, in addition to sampling loss due to warming of the fluid was possible. Aluminum foil was chosen as the initial intent was to perform all sampling with UF probes in addition to direct sampling and using snap-on or screw-on lids would have damaged the probes. The study was converted to direct sampling only at the last minute due to concerns of probe functioning and continued patency during the initial assembly, and fluid was already in the dishes being prewarmed. The evaporative fluid loss could account for the increase in Ag concentration of the fluid at the end of the study. No attempt was made to estimate fluid losses during the study, as measuring at each time point would have necessitated interrupting the elution and manipulating the specimen and might cause additional fluid loss. However, measuring the volume at 96 hours could have been performed. Fluid loss during a continuous release elution model is a known limitation of the model chosen [24], however removing and replacing all fluid at each time point might artificially keep a larger gradient intact than a continuous model would. In addition, a continuous model might more closely resemble the *in vivo* decrease in gradient between the slow-release compound and its environment.

The concentration of Ag measured in the samples obtained both by UF probe and pipette sampling decreased over time. This could be explained by the stagnant nature of the fluids within the tube, although care was taken to sample from the lowest area of the fluid in case of sedimentation. Elution experiments ideally are performed with constant stirring [24], or agitation prior to sampling [16]. We opted to not agitate the tubes in order to keep the probes fully submerged at all times. In addition, the probe would remain in the same place *in vivo* as well, and a stationary fluid was felt to be appropriate given that no concentration gradient was present. Pipette samples were obtained from the area in the tube in the location of filtration membranes of the UF as to not introduce a variable between the two methods. The higher variation seen in samples obtained by UF probes is interesting and indicates that care must be taken with *in vivo* sampling to account for variability, for example by taking multiple samples at each time point.

Additional limitations could be inconsistency in mixing the specimens and solutions as well as sampling or measurement errors. We included 3 repeats of the elution study to minimize the effect of both inconsistency and sampling. Six repeat samples were used in the static sampling study, however inconsistency between the solutions could still exist.

## Conclusions

Elution into plasma and UF probe sampling of Ag in plasma are possible. While use of DPBS might be more cost effective, plasma should be considered for *in vitro* silver elution studies due to difference in elution and recovery. Silver nanoparticles in poloxamer 407 might release more slowly than other drugs that have been studied under similar conditions. Both UF probe sampling and direct pipette sampling underestimate the actual Ag concentration in the fluid. Ultrafiltration probe sampling of Ag is possible and UF probes could be used to measure *in vivo* local tissue concentrations, but the possibility of variations in the sampled fluid should be kept in mind.

## Supporting information

Supplemental file - original data

## Acknowledgements

The authors thank the Nanomedicines Characterization Core Facility (NCore) at the Center for Nanotechnology in Drug Delivery (CNDD) and Dr Marina Sokolsky for their assistance with sample and data analysis.

